# Old Meets New: Combining Herbarium Databases with Genetic Methods to Evaluate the Invasion Status of Baby’s Breath (*Gypsophila paniculata*) in North America

**DOI:** 10.1101/686691

**Authors:** Sarah K. Lamar, Charlyn G. Partridge

## Abstract

**Aim:** This paper aims to inform our knowledge of common baby’s breath’s (*Gypsophila paniculata*) current population structure and invasion status using a combination of contemporary genetic methods and historical herbarium data.

**Taxon:** *Gypsophila paniculata* (Angiosperms: Eudicot, Caryophyllaceae)

**Location:** Samples were collected from seven locations spanning a portion of the plant’s North American range: Washington, North Dakota, Minnesota, and Michigan, United States.

**Methods:** To analyze contemporary population structure, individuals of *G. paniculata* from 7 distinct sampling locations were collected and genotyped at 14 microsatellite loci. Population structure was inferred using both Bayesian and multivariate methods. To investigate *G. paniculata’*s invasion status, public herbarium databases were searched for mention of the species. Records were combined, resulting in a database of 307 herbarium collections dating from the late 1800’s to current day. Using this database, invasion curves were created at different geospatial scales.

**Results:** Results of genetic analyses suggest the presence of at least two genetic clusters spanning our seven sampling locations. Sampling locations in Washington, North Dakota, Minnesota, and northwestern Michigan form one genetic cluster, distinct from our two more southern sampling locations in Michigan, which form a second cluster with increased relative genetic diversity. Invasion curves created for these two clusters show different time periods of invasion. An invasion curve created for North America suggests *G. paniculata’*s range may still be expanding.

**Main conclusions:** *Gypsophila paniculata* has likely undergone at least two distinct invasions in North America, and its range may still be expanding. Restricted genetic diversity seen across a wide geographic area could be a signature of limited seed distributors present during the early period of this garden ornamental’s invasion.

## Introduction

Biological invasions are a growing concern in the era of global trade and transport. In the United States alone, there have been over 50,000 introductions of plant, animal, and microbe species into environments beyond their native range (Pimentel, Zuniga, & Morrison, 2005). These introductions can have dramatic impacts on native flora and fauna; roughly 42% of species listed on the Endangered Species Act are threatened by competition with invasives (Wilcove, Rothstein, Dubow, Phillips, & Losos, 1998). Of particular concern among invasive species are invasive weeds, a group that currently spreads across the United States at a rate of 700,000 ha/year (Pimentel et al., 2005). This rapid consumption of land by non-native species makes managing invasive weeds a priority for the preservation of native ecosystems and the native biota that inhabit them.

Many plant and animal species that are transported into new environments will not become problematic invaders, defined as species not native to an area whose range or abundance is increasing regardless of habitat (P. Pyšek, 1995; Williamson & Fitter, 1996). Non-native species that go on to become invasive in their new environments must survive transport, reproduce as a relatively small founding population, respond to potentially novel environmental stressors, and overcome the “lag phase” of an invasion (Larkin, 2012; Williamson & Fitter, 1996). This lag phase is characterized by a period of slow growth after initial introduction that, if overcome, can lead to a period of rapid population expansion before eventually plateauing as the new range is saturated (Mack et al., 2000). Despite the many barriers that species face on the road to becoming invasive, the impacts of these events are a growing cause for concern.

As the number of global invasion events increases, so does the importance of developing and implementing cost effective methods for studying invasion events. Invasion curves are one such tool used to assess an invasive species’ status and rate of spread (see Antunes & Schamp, 2017; Shih & Finkelstein, 2008). Invasion curves can offer researchers important insight to a species’ lag time after introduction into new environments, providing valuable information associated with response time, geographic barriers to spread, and the efficacy of existing management strategies (Antunes & Schamp, 2017; Crooks, 2007). Because they are crafted using historical data, such as herbarium records, invasion curves are both cost effective and capable of offering important glimpses into the often-unnoticed lag phase of an invasion (Antunes & Schamp, 2017). Invasion curves have been used to recognize potential refuges for weed species (e.g. Lavoie, Jodoin, & De Merlis, 2007), identify major drivers of invasive species spread (e.g. Fuentes, Ugarte, Kühn, & Klotz, 2008; Petr Pyšek, Jarošík, Müllerová, Pergl, & Wild, 2008), and even help assess the efficacy of potential biocontrol agents (e.g. Boag & Eckert, 2013).

While invasion curves are useful for addressing many questions managers and researchers may have, they are limited by the constraints associated with herbarium records and survey data. To supplement these constraints, genetic analyses may be used to provide information concerning contemporary gene flow, adaptive potential, relatedness among invasive populations, and possible resistance to control efforts (e.g. Abdelkrim, Pascal, Calmet, & Samadi, 2005; Zalewski et al., 2010). Genetic analyses of invasive species has been used to identify potential barriers to migration (Haynes, Gilligan, Grewe, & Nicholas, 2009) and estimate the number of likely invasion events a species may have undergone (Meimberg et al., 2010) While this information can help improve our understanding of invasive science as a whole, it also has immediate benefits to managers; because distinct genetic populations have different potential evolutionary trajectories, understanding the genetic structure of populations is critical for effective management (Moritz, 1994; Palsbøll, Bérubé, & Allendorf, 2007).

*Gypsophila paniculata* (common baby’s breath) is a perennial forb native to the Eurasian steppe region (Darwent, 1975; Darwent & Coupland, 1966). *Gypsophila paniculata* is characterized by a taproot that can reach several meters deep, which is thought to help the plant to out-compete natives for limited resources in harsh environments (Darwent & Coupland, 1966). Though it does not produce floral primordia until at least its second year, *G. paniculata* can yield almost 14,000 seeds per growing season (Darwent & Coupland, 1966; Stevens, 1957). These seeds are small (86mg/100 seeds) and primarily distributed by wind forces; when plants reach senescence, they break off above the caudex and form tumbleweeds that spread seeds as they roll (Darwent & Coupland, 1966; Stevens, 1957).

Populations of *G. paniculata* were established in North America by the late 1880’s, likely having been introduced due to its popularity in the garden and floral industries (Darwent & Coupland, 1966). According to the Early Detection and Distribution Mapping System, *G. paniculata* can now be found growing as an invasive species in 30 U.S. states (EDDMapS, 2019). It has been listed as a Class C (widespread noxious weed) in Washington and California and is considered a priority invasive by Michigan Department of Natural Resources (Emery & Doran, 2013; Michigan Department of Natural Resources, 2015; Swearingen & Bargeron, 2016). *Gypsophila paniculata* can form dense stands in the areas that it invades; in some parts of Sleeping Bear Dunes National Lakeshore, an invaded area in Michigan, *G. paniculata* forms as much as 75% of the vegetation present (Karamanski, 2000; Rice, 2018). These dense monocultures can have impacts on native plant, nematode, and arthropod communities, potentially having ripple effects across the trophic system (Emery & Doran, 2013; Reid & Emery, 2018).

To help understand the invasion status of this problematic plant species, this study aims (1) to define the population structure of contemporary *G. paniculata* growing throughout a portion of its introduced range, and (2) to create invasion curves of *G. paniculata* to assess its current invasion status at different geospatial scales.

## Methods

### Study Sites and Contemporary Sample Collection

To investigate contemporary population structure of *G. paniculata,* tissue samples from five locations across the United States were collected in the summer of 2018: Petoskey, MI; Knife River Indian Villages National Historic Site, ND; Ottertail, MN; Chelan, WA; and Osborne Bay, WA (Figure 1, Table 1). Samples from two additional locations in Sleeping Bear Dunes National Lakeshore, MI and Arcadia Dunes, MI were collected in the summer of 2016 (Table 1) (Leimbach-Maus, Parks, & Partridge, 2018a). Leaf tissue was collected from 15-30 individuals per location (5-10 leaves per plant). Tissue samples were placed inside coin envelopes and stored in silica until DNA extraction. Individuals were collected for sampling by identifying a plant of any size separated from other sampled individuals by at least 2 meters, in efforts to minimize the likelihood of sampling closely related plants.

**Figure 1.**
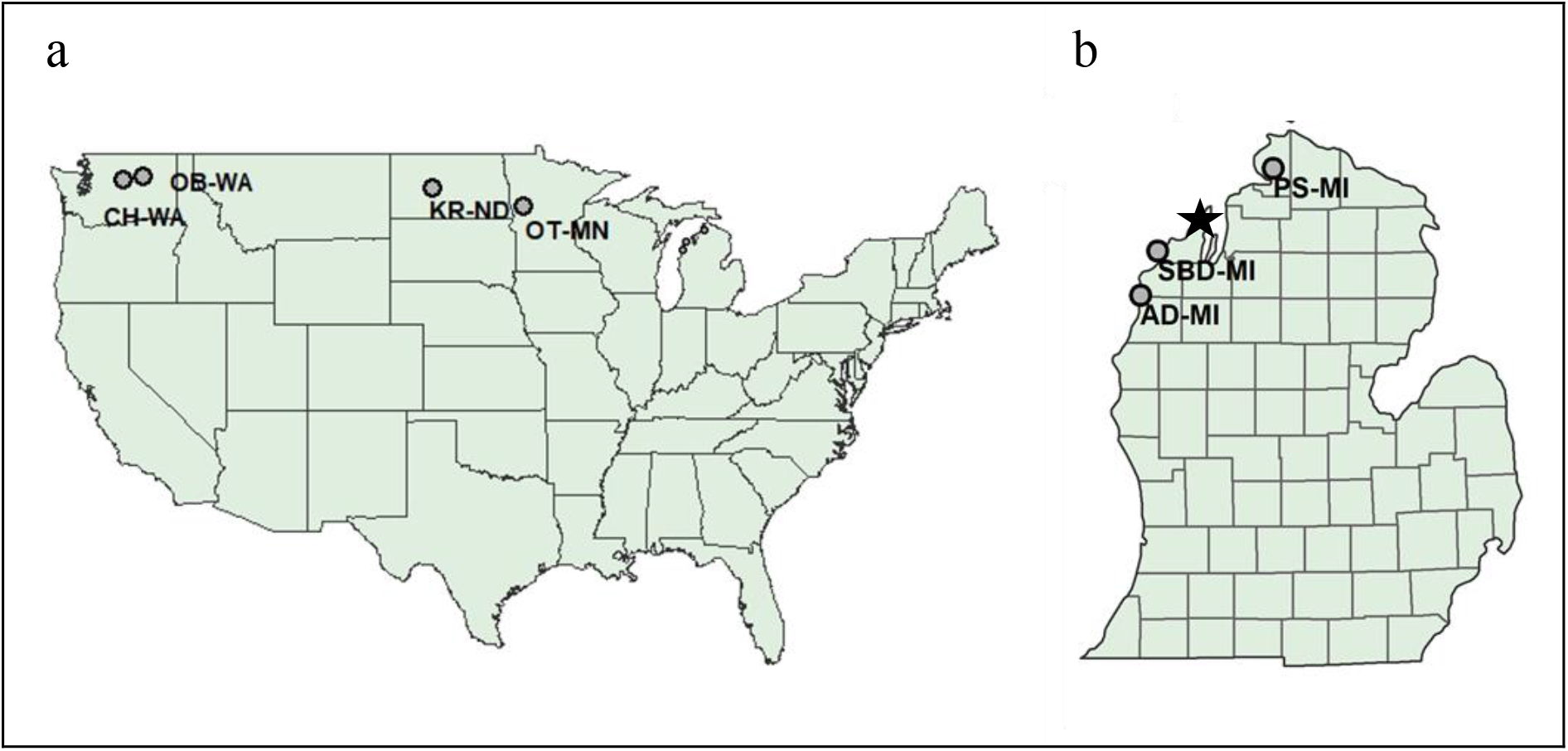
Sampling locations for assessing *G. paniculata* population structure used in this study; locations in Washington, North Dakota, and Minnesota are visualized in panel (a), locations in Michigan are visualized in panel (b). The Leelanau Peninsula is denoted by a black star. Sampling location codes: Chelan, WA (CH-WA); Osborne Bay, WA (OB-WA); Knife River Historic Indian Villages, ND (KR-ND); Ottertail, MN (OT-MN); Petoskey State Park, MI (PS-MI); Sleeping Bear Dunes National Lakeshore, MI (SBD-MI); Arcadia Dunes, MI (AD-MI).

**Table 1.**
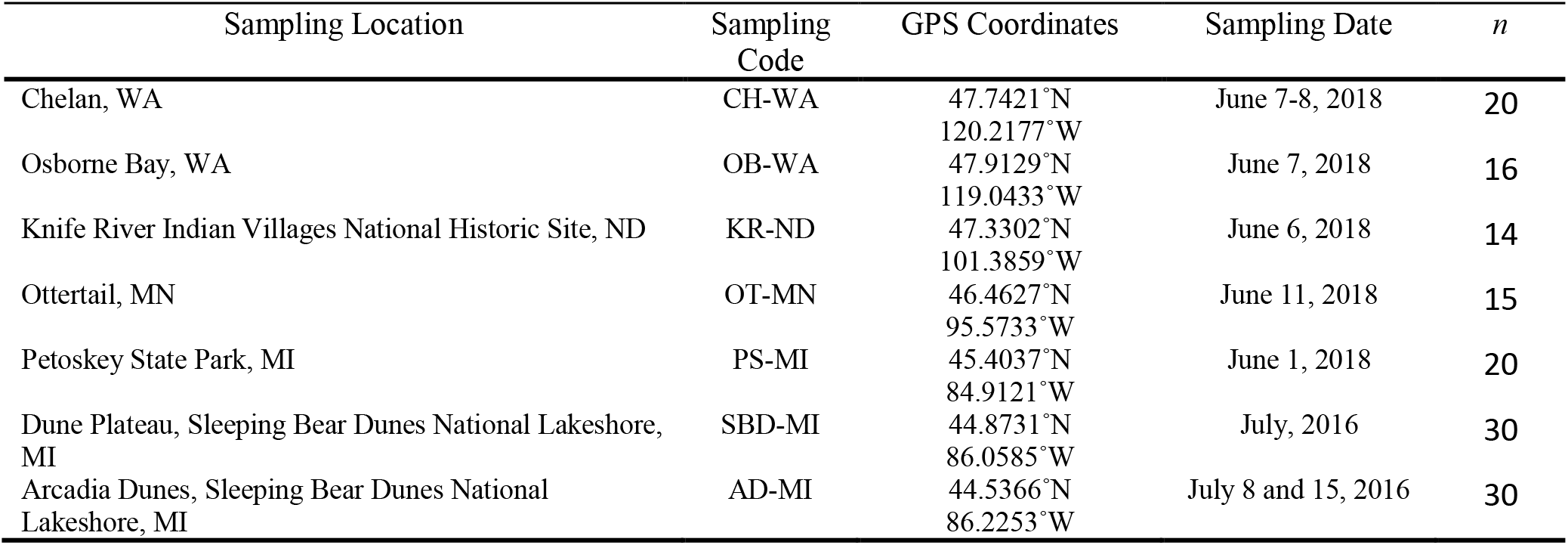
Locations, dates, sample size, and geographic coordinates for analyzed samples of baby’s breath (G. *paniculata*).

### Microsatellite Analysis

For each sample (n=145), 0.25 g of dried leaf tissue was weighed out. DNA was extracted from tissue samples using a Qiagen DNeasy plant mini kit (QIAGEN, Hilde, Germany); manufacturer instructions were largely followed, apart from an extra wash with AW2 buffer. Extracted DNA was run through a Zymo OneStep PCR Inhibitor Removal Column (Zymo, Irvine, CA) twice.

Samples were amplified at 14 nuclear microsatellite loci identified as polymorphic and specific to *G. paniculata* (Leimbach-Maus, Parks, & Partridge, 2018b) (Table S1). PCR was conducted using a 5’ fluorescently-labelled primer (6-FAM, PET, NED, or VIC) (Applied Biosystems, Foster City, CA) and an unlabeled reverse primer. Reaction mixtures consisted of 1x KCl buffer, 2.0-2.5 mM MgCl_2_, 300 μM dNTP, 0.08 mg/mL BSA, 0.4 μM forward primer, 0.4 μM reverse primer, 0.25 units Taq polymerase, and 50 ng DNA template. The thermal cycling profile consisted of 5 minutes of denaturation at 94°C, followed by 35 cycles of 94°C for 1 minute, 1 minute of annealing at 62° (with the exception of locus BB_2888, see Table S1), 1 minute of extension at 72°C, and a final elongation step of 10 minutes at 72°C. PCR products were visualized on a 2% agarose gel using GelRed™ (Biotium, Freemont, CA) before multiplexing with consideration to dye color and allele size (Table S1). Genescan 500 LIZ size standard (Thermo Fisher Scientific, Waltham, MA) was added to multiplexed product with Hi-Di™ Formamide (Thermo Fisher Scientific, Waltham, MA) to aid in denaturing. Fragment analysis was conducted on an ABI3130xl Genetic Analyzer (Applied Biosystems, Foster City, CA). Individuals were genotyped using the automatic binning procedure on GENEMAPPER v5 (Applied Biosystems, Foster City, CA) before being visually verified to reduce error. A subsample of 20 individuals were genotyped twice to ensure consistent allele scoring.

### Exploratory Data Analysis

The presence of null alleles was investigated using MICRO-CHECKER v2.2.3; using this method, none were found (Van Oosterhout, Hutchinson, Wills, & Shipley, 2004). Data were screened using the ‘STRATAG’ package in the R statistical program v3.4.3 (Archer, Adams, & Schneiders, 2016; R Development Core Team, 2017) for any individual that was missing greater than 20% of loci and any locus that was missing greater than 10% of individuals; on this basis, no data were removed.

### Measures of Genetic Diversity

Linkage disequilibrium and a test for Hardy-Weinberg equilibrium were calculated using GENEPOP v4.6 with 1,000 batches of 1,000 Markov chain Monte Carlo iterations (Raymond & Rousset, 1995; Rousset, 2008). There was no significant deviation from linkage equilibrium across populations and no data were removed on this basis. Expected versus observed heterozygosity, number of private alleles, and Weir and Cockerham’s population pairwise F_ST_ values were conducted using GENALEX v6.502 in Microsoft Excel (Peakall & Smouse, 2006, 2012; Weir & Cockerham, 1983). Inbreeding coefficient (F_IS_) values were calculated in GENEPOP.

### Genetic Structure

A principal coordinate analysis (PCoA) was conducted using a genetic distance matrix in GENALEX (Peakall & Smouse, 2006, 2012). Population clustering was analyzed in STRUCTURE v2.3.2(Pritchard, Stephens, & Donnelly, 2000) using an admixture model, both with and without *a priori* location information, and a burn-in length of 100,000 with 1,000,000 MCMC replicates after burn-in. Ten iterations were run for each *K* value (1-9). The number of genetic clusters was determined using the Evanno ΔK method (Evanno, Regnaut, & Goudet, 2005). Because ΔK is based on a rate of change, it does not evaluate K=1 and can be biased towards K=2 (Dupuis et al., 2017). Considering this, we also used discriminant analysis of principal components (DAPC) to support our STRUCTURE findings (Jombart, Devillard, & Balloux, 2010). DAPC separates variance into within-group and between-group categories and works to maximize cluster discrimination; this analysis was conducted using the package ‘adegenet’ v2.1.1 in R (Jombart et al., 2010). Because retaining too many principal components (PC’s) can lead to instability in cluster membership properties, a cross-validation was performed to inform the analysis of the optimal number of PC’s. After cross-validation, 16 of 28 PC’s and all eigenvalues were retained. An analysis of molecular variance (AMOVA) was run using 9,999 permutations in GENALEX to test how much variance could be explained by between-population and within-population variation.

### Herbarium Invasion Curves

To create invasion curves for *G. paniculata* population clusters, public herbarium databases were searched for records of this species; species identification was visually confirmed when possible. Records that did not include location data (either GPS, county (U.S.) or regional municipality (Canada)) and year were discarded, resulting in 307 records from 65 North American institutions. All locality information was standardized to the county scale to reduce the risk of redundant specimen collection while maintaining adequate resolution (Antunes & Schamp, 2017). Earliest samples were found in the late 1890’s-early 1900s in California, Michigan, Minnesota, and New York and this is consistent with the earliest times in which *G. paniculata* seeds were first being sold in the United States (1886), based on a search of the Henry G. Gilbert Nursery and Seed Trade Catalog Collection from the Biodiversity Heritage Library (https://www.biodiversitylibrary.org/).

To examine the invasion status of populations belonging to genetic clusters identified from our population genetics analysis, herbarium records were grouped according to desired geospatial scales (cumulative North America, genetic cluster 1, and genetic cluster 2). Only records for the first occurrence of *G. paniculata* in each county or regional municipality were kept. Cumulative records for North America had 184 unique municipalities represented, while both genetic clusters had fewer unique localities (cluster 1 = 42, cluster 2 =16) and required log transformation for better visualization. Data were plotted as the cumulative number of localities invaded over time using the statistical program R.

## Results

### Measures of Genetic Diversity

Overall, the five western populations and northernmost Michigan population showed lower levels of genetic diversity compared with the two more southern populations in Michigan (Table 2). Pairwise comparisons yielded significant F_ST_ values between all populations; however, SBD-MI, AD-MI, and KR-ND showed comparatively high pairwise F_ST_ values compared to other populations (Table 3). F_ST_ values between CH-WA, OB-WA, and PS-MI were relatively low compared to other sample locations in this study, suggesting more limited genetic differentiation among these populations (Table 3).

**Table 2.**
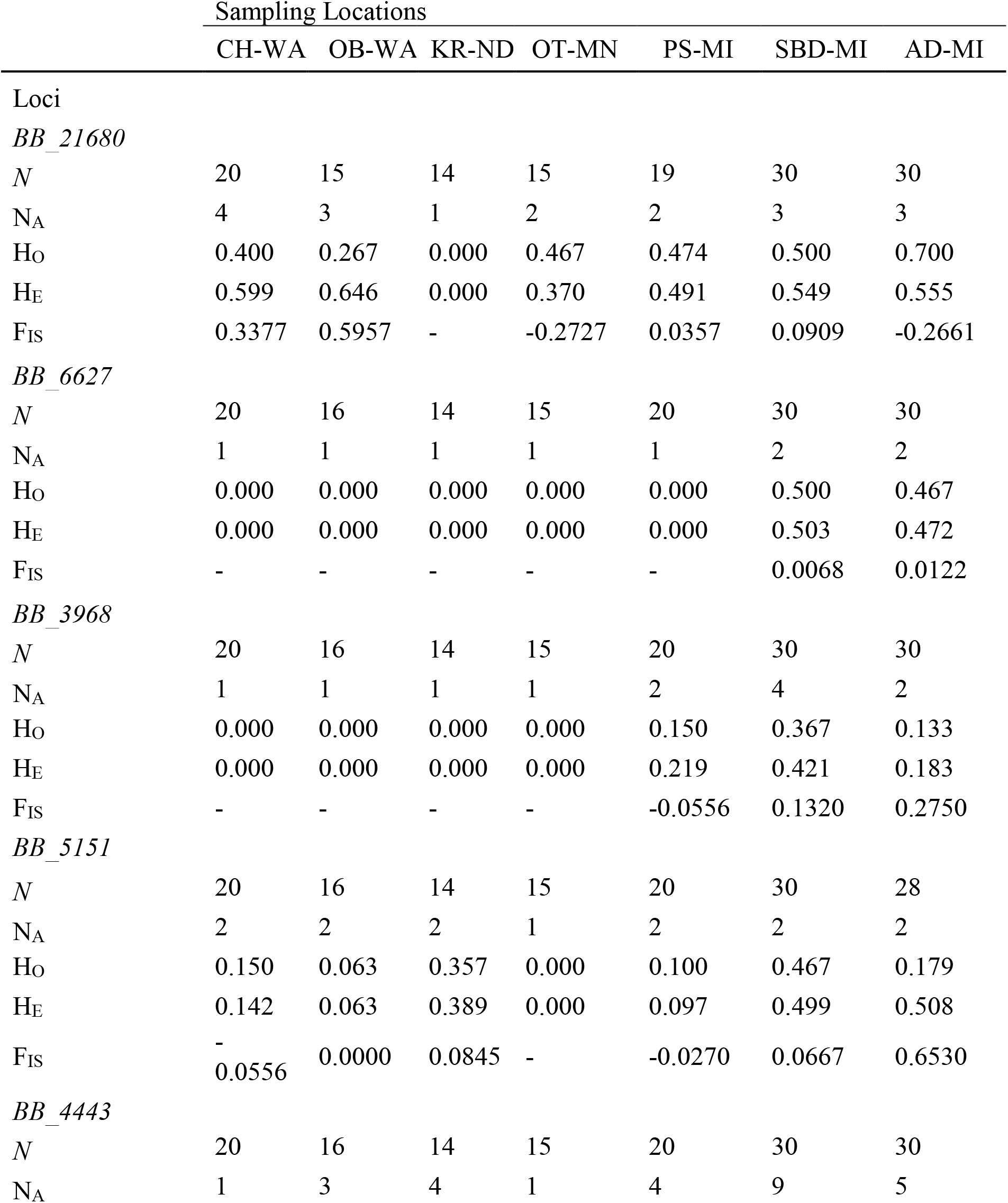
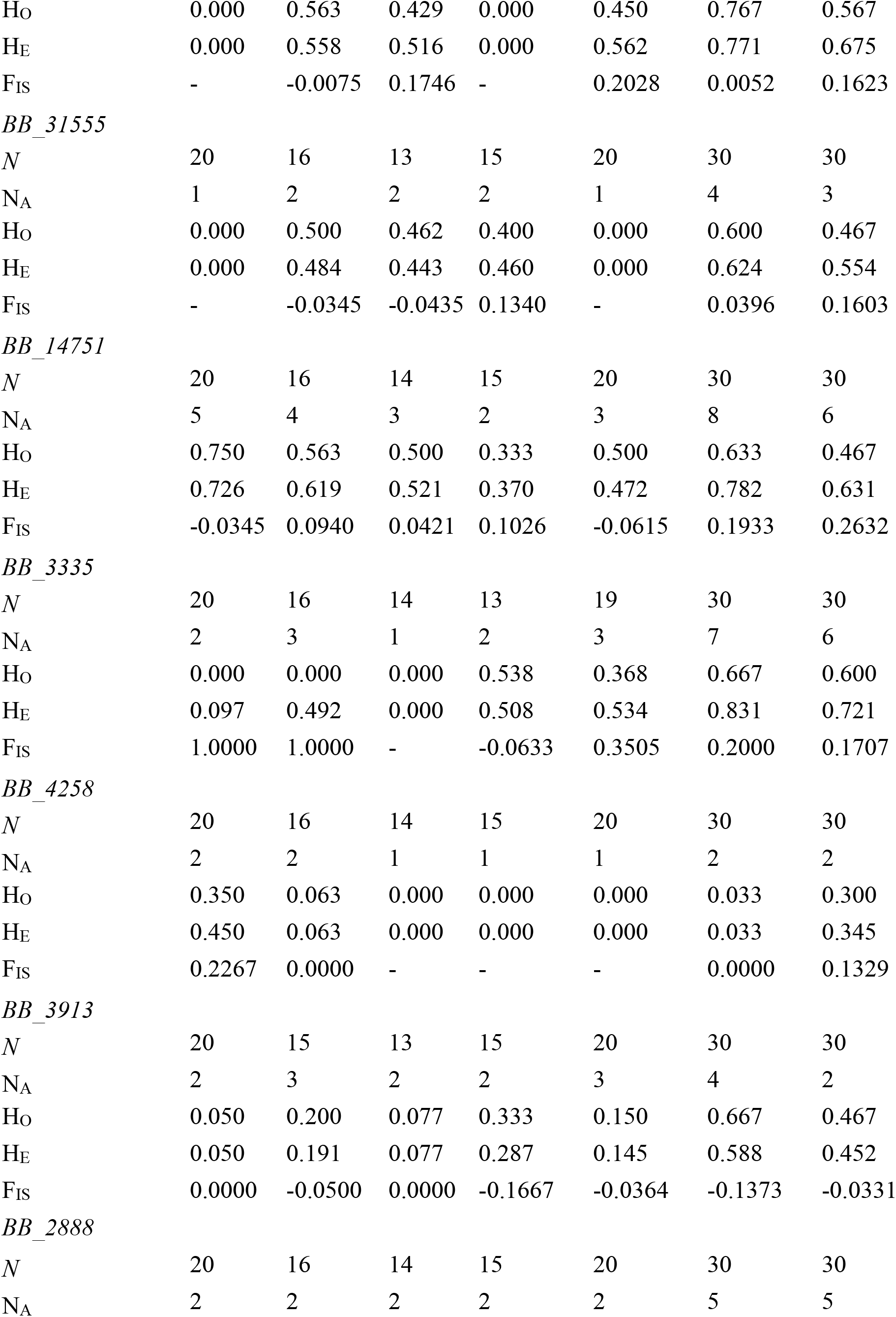
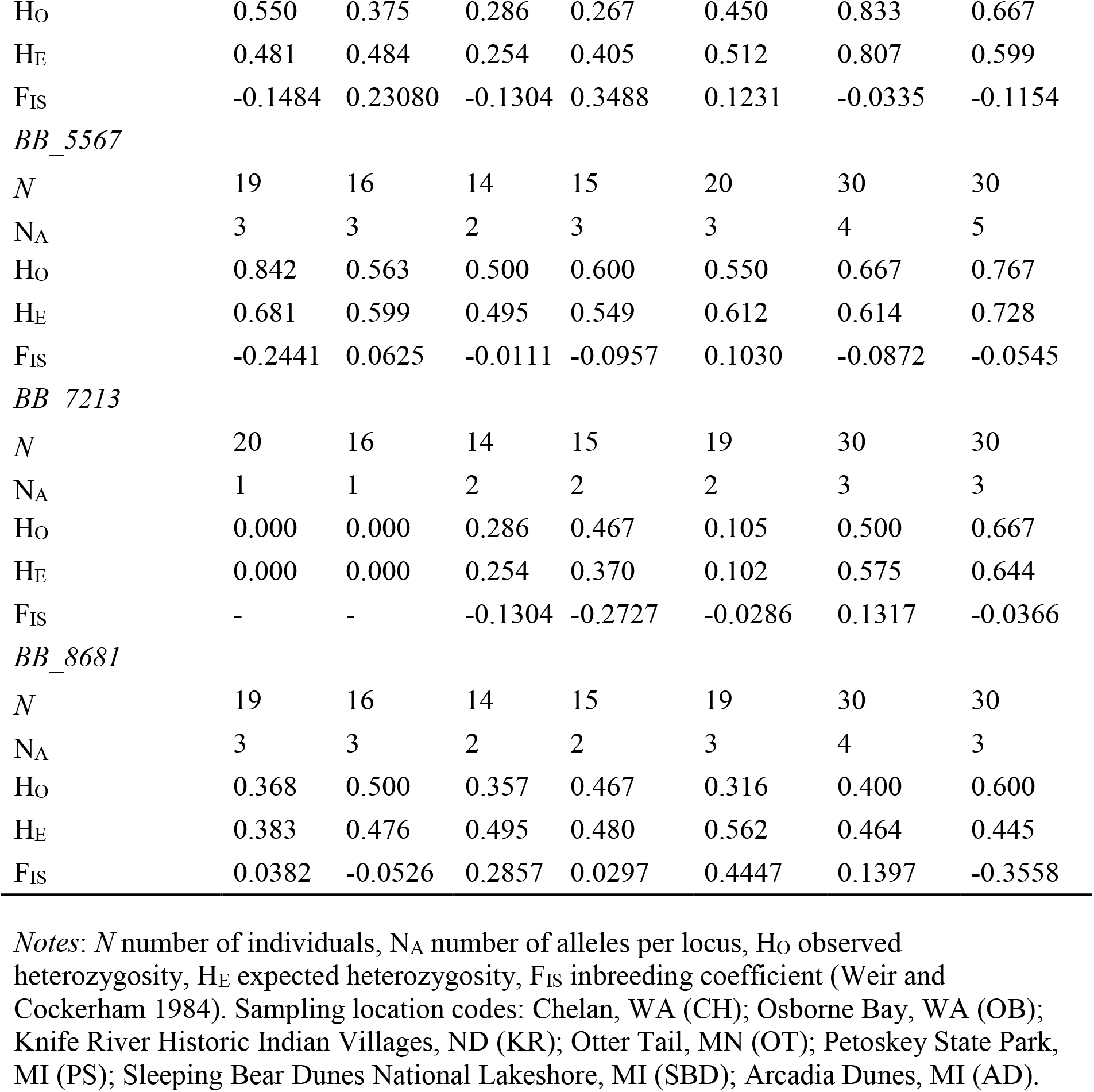
Genetic diversity measures for seven *G. paniculata* sampling locations sequenced at 14 microsatellite (nSSR) loci.

**Table 3.** Population pairwise F_ST_ (Weir and Cockerham, 1984) for *G. paniculata* populations using microsatellite data calculated in GenAlEx 6.502 (Peakall and Smouse, 2006, 2012) running 9,999 permutations. Darker colors indicate increasing (higher) values; all values are significant with p-values <0.05. Sampling location codes: Chelan, WA (CH-WA); Osborne Bay, WA (OB-WA); Knife River Historic Indian Villages, ND (KR-ND); Ottertail, MN (OT-MN); Petoskey State Park, MI (PS-MI); Sleeping Bear Dunes National Lakeshore, MI (SBD-MI); Arcadia Dunes, MI (AD-MI).

|  | CH-WA | OB-WA | KR-ND | OT-MN | PS-MI | SBD-MI | AD-MI |
| --- | --- | --- | --- | --- | --- | --- | --- |
| CH-WA | – |  |  |  |  |  |  |
| OB-WA | 0.077 | – |  |  |  |  |  |
| KR-ND | 0.188 | 0.141 | – |  |  |  |  |
| OT-MN | 0.124 | 0.104 | 0.194 | – |  |  |  |
| PS-MI | 0.111 | 0.075 | 0.150 | 0.094 | – |  |  |
| SBD-MI | 0.202 | 0.131 | 0.201 | 0.196 | 0.173 | – |  |
| AD-MI | 0.192 | 0.153 | 0.188 | 0.168 | 0.160 | 0.070 | – |

### Genetic Population Structure

Results of the Bayesian clustering analysis conducted in the program STRUCTURE suggest two population clusters (K=2), both from ΔK and Ln Pr (X|K) (Figure S1). Analysis was conducted both with and without prior sampling location; there was no observable difference between the two (without priors shown in Figure 2). Cluster 1 is comprised of sampling locations in North Dakota, Minnesota, Washington, and the northernmost site in Michigan; cluster 2 is comprised of the two more southern sites in Michigan (Figure 2). Overall, there is little admixture between the two groupings, with only few individuals in AD-MI showing any signs of genetic mixing.

**Figure 2.**
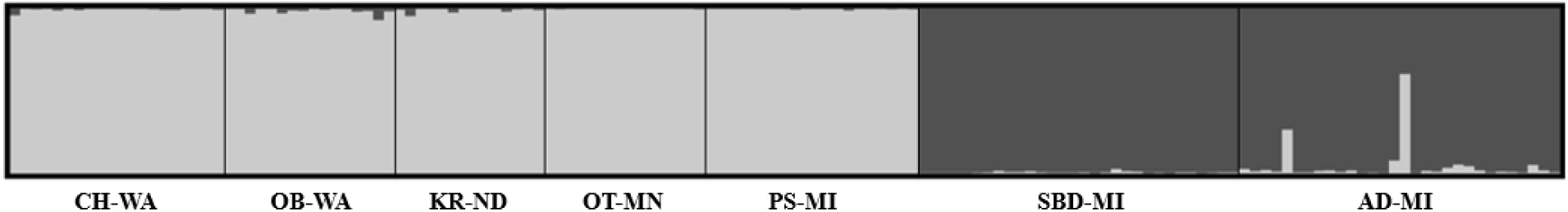
Results of Bayesian cluster analysis of *G. paniculata* genotyped at 14 microsatellite loci, performed using the program STRUCTURE (Pritchard et al. 2000). Each individual (n=145) is represented by a single column, with different colors indicating the likelihood of assignment to that cluster. Black lines delineate sampling location. Results suggest 2 population clusters (K=2). Locations are listed from west to east and north to south (MI). Sampling location codes: Chelan, WA (CH-WA); Osborne Bay, WA (OB-WA); Knife River Historic Indian Villages, ND (KR-ND); Ottertail, MN (OT-MN); Petoskey State Park, MI (PS-MI); Sleeping Bear Dunes National Lakeshore, MI (SBD-MI); Arcadia Dunes, MI (AD-MI).

Population structure was further analyzed with a PCoA based on a genotypic distance matrix. Population division along the primary principal coordinate accounted for 27.22% of variation present. Along this coordinate, the trends seen in STRUCTURE analysis were supported, with populations SBD-MI and AD-MI separating out from the remaining five populations (Figure 3). The secondary principal component suggests further separation may exist between SBD-MI and AD-MI (9.80% of variation present) if *K* is forced to 3. The grouping of CH-WA, OB-WA, OT-MN, KR-ND, and PS-MI into the same cluster is supported by this analysis.

**Figure 3.**
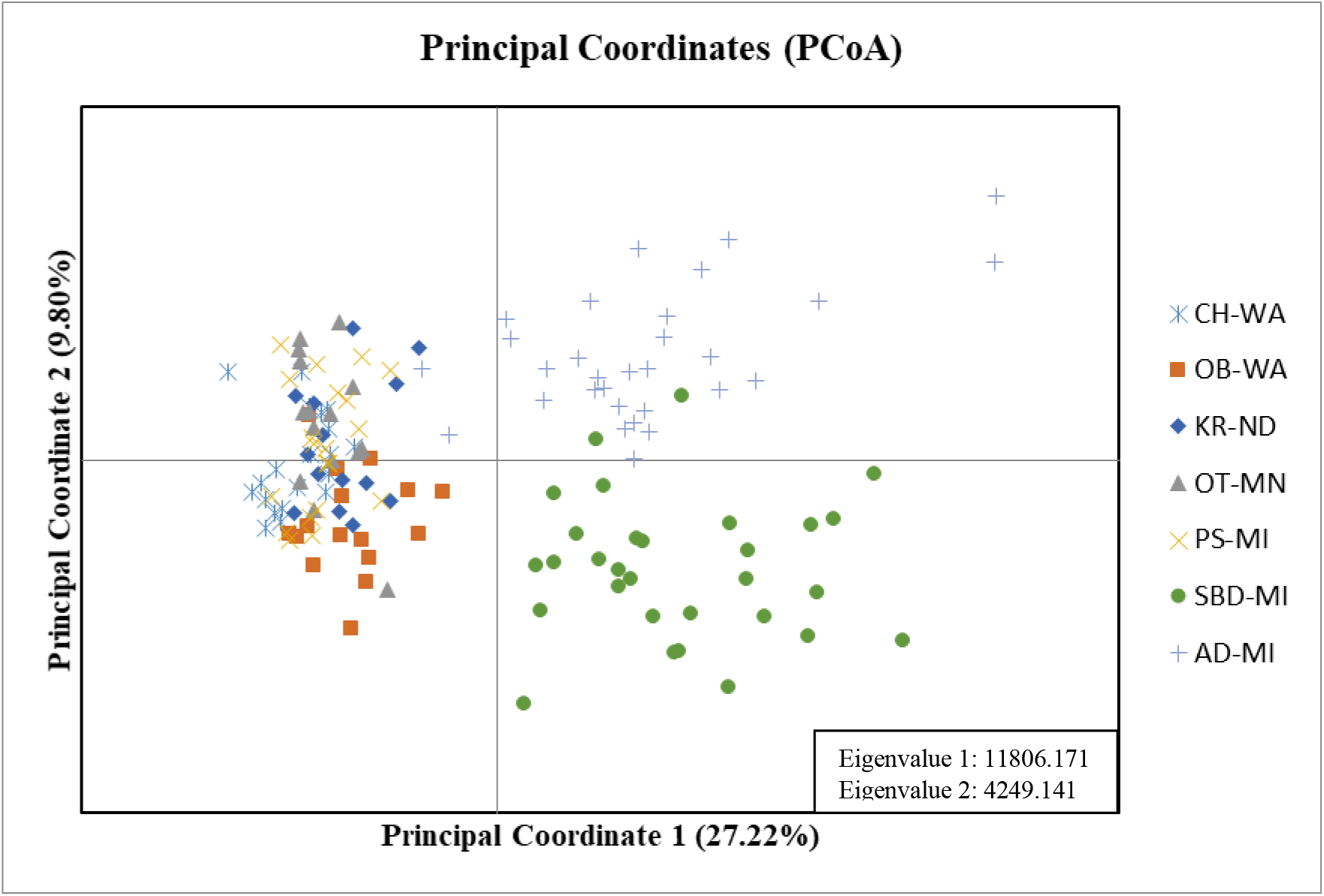
Principal Coordinates Analysis (PCoA) of seven geographic populations of baby’s breath *(G. paniculata*) genotyped at 14 microsatellite loci, based on a genotypic distance matrix, and performed in GenAlEx 6.502 (Peakall and Smouse, 2006,2012). Sampling location codes: Chelan, WA (CH-WA); Osborne Bay, WA (OB-WA); Knife River Historic Indian Villages, ND (KR-ND); Ottertail, MN (OT-MN); Petoskey State Park, MI (PS-MI); Sleeping Bear Dunes National Lakeshore, MI (SBD-MI); Arcadia Dunes, MI (AD-MI).

DAPC’s Bayesian Information Criterion suggested either 2 or 3 genetic clusters (Figure S2). Sampling locations in Arcadia Dunes, MI and Sleeping Bear Dunes, MI separated into distinct populations when *K* was pushed to 3, in order to investigate all cluster possibilities (Figure 4a). Individual membership to clusters is detailed in Figure 4c, which shows that cluster 1 is 82% comprised of individuals from SBD-MI, cluster 2 is 93% comprised of individuals from AD-MI, and cluster 3 has a relatively even contribution of individuals from CH-WA, OB-WA, KR-ND, OT-MN, and PS-MI. When individual distribution is viewed along the primary discriminant function, overlap between clusters 1 (SBD-MI) and 2 (AD-MI) is clearly visible (Figure 4b), while cluster 3 shows little to no overlap with clusters 1 or 2.

**Figure 4.**
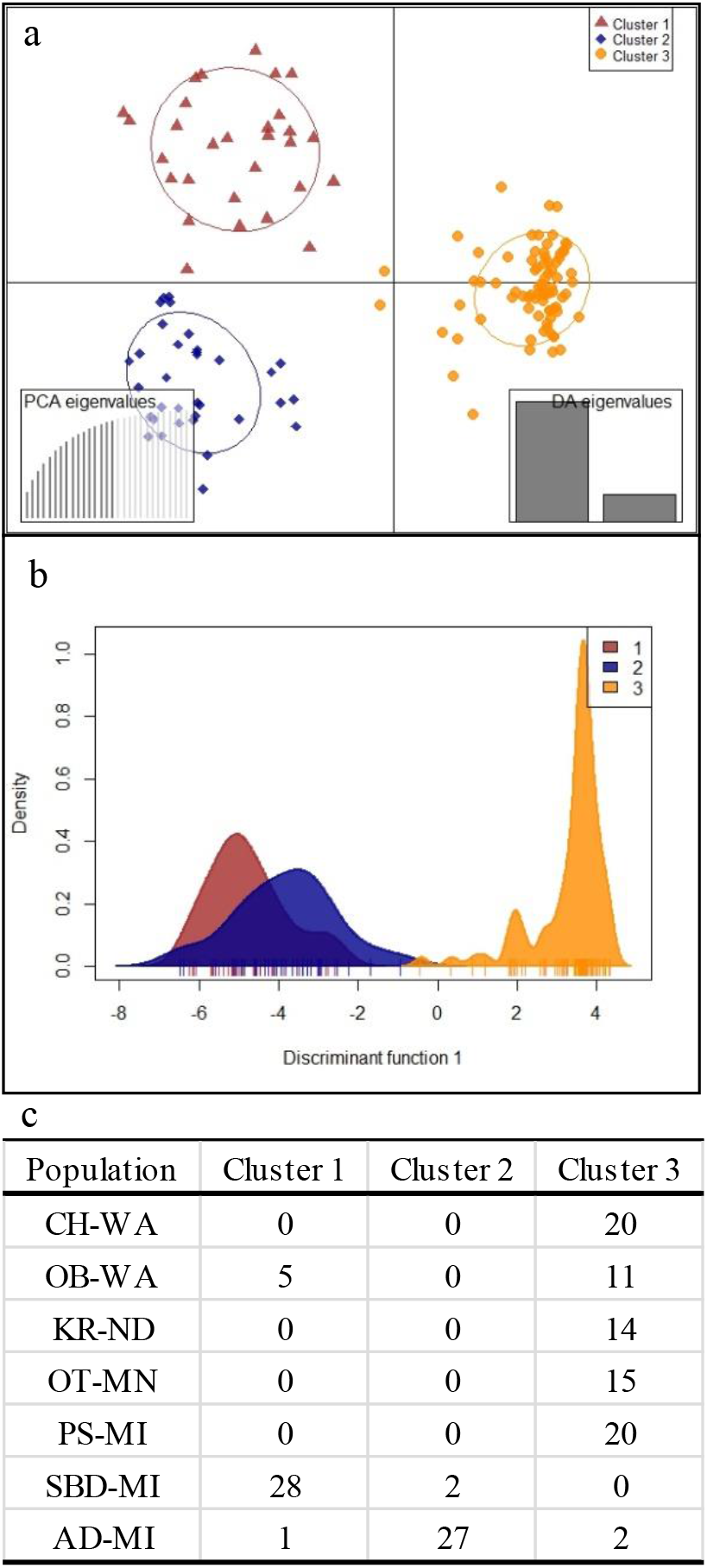
Discriminant analysis of principal components (DAPC) based on *G. paniculata* analyzed at 14 microsatellite loci and calculated in the *‘adegenet’* package for R (Jombart et al., 2010). (a) Scatterplot showing both discriminant function axes and eigenvalues. Each point represents an individual (n=145). After cross validation, 16 of 28 PC’s were retained. (b) Plot visualizing DAPC sample distribution on the primary discriminant function. (c) Individual assignment to clusters using all eigenvalues explained by the PCA. Sampling location codes: Chelan, WA (CH-WA); Osborne Bay, WA (OB-WA); Knife River Historic Indian Villages, ND (KR-ND); Ottertail, MN (OT-MN); Petoskey State Park, MI (PS-MI); Sleeping Bear Dunes National Lakeshore, MI (SBD-MI); Arcadia Dunes, MI (AD-MI).

AMOVA results show that a significant amount of variation could be explained by differences among populations within regions (Φ_PR_= 0.229, p <0.001) and by differences between our first region (CH-WA, OB-WA, KR-ND, OT-MN, PS-MI) and second region (SBD-MI and AD-MI) (Φ_RT_ = 0.246, p <0.001). However, most variation present was found within populations (Φ_PT_ = 0.419, p<0.001).

### Invasion Curves

Invasion curves created using herbarium records, standardized to the scale of local municipality, were used to visualize the invasion stage (i.e. lag phase, expansion phase, or plateau phase) of *G. paniculata* at various geospatial scales (Figure 5). Records for North America slowly accumulate during the early periods of invasion (1890’s) until roughly the 1940’s, after which the number of records being collected in new localities begin to accumulate rapidly (Figure 5b). This likely represents the shift from the initial lag phase of invasion to the expansion phase. With no clear plateau being reached, the expansion phase of *G. paniculata* across the entirety of North America appears to continue. Considering records according to assignment with genetic cluster, initial collection for cluster 1 (WA, ND, MN, and PS-MI) is noted in the late 1890’s, but few additional records were archived until the mid-1920’s, when herbarium data for *G. paniculata* suggest an expansion of this population (Figure 5c). A plateau can be seen beginning in the mid 1990’s when the curve of the line begins to taper. Records for genetic cluster 2 (Figure 5d) are comprised of collections from mid-southwest Michigan (defined as south of the Leelanau Peninsula, based on results from this study and a previous study conducted by Leimbach-Maus et al., 2018a). Rapid expansion began shortly after its first collection in the late 1940’s, with the spread beginning to plateau around 1970. No discernable lag period is noted in the collection data for this cluster.

**Figure 5.**
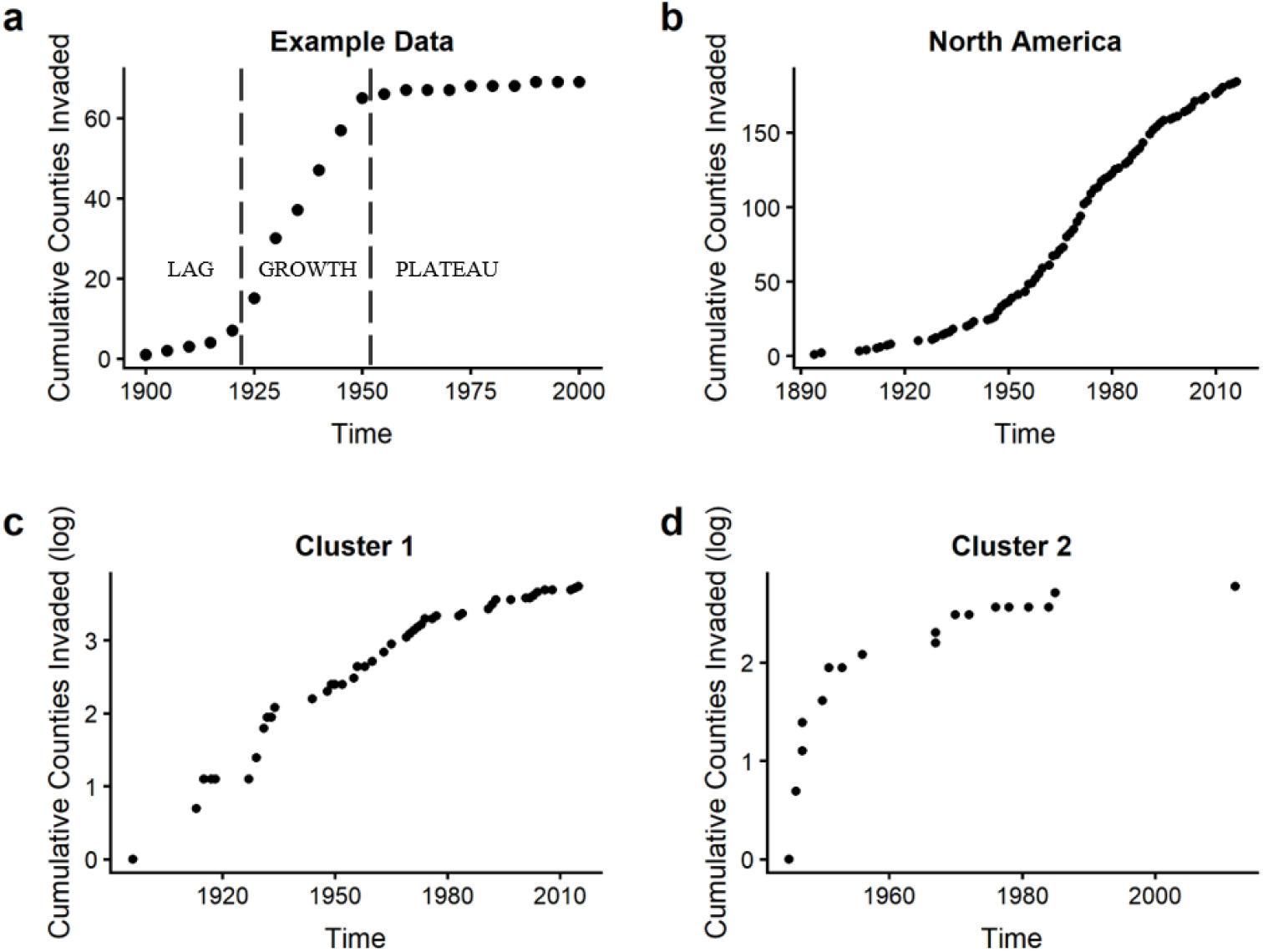
Invasion curves created using herbarium data for *Gypsophila paniculata* collection in (b) North America, (c) genetic cluster 1, and (d) genetic cluster 2 (a gap in sample collection is evidence by the lack of points on the graph). An example invasion curve illustrating the three-stage invasion pathway typical of many invasions is visualized in panel (a). Cluster assignment: (1) Washington, North Dakota, Minnesota, and northwest Michigan. (2) Michigan south of the Leelanau Peninsula.

## Discussion

Our data from populations of *G. paniculata* growing across a portion of its introduced range in North American reveals the presence of at least two distinct genetic clusters. The northernmost sampling location in Michigan (PS-MI) clustered with the four sampling locations located across North Dakota, Minnesota, and Washington, and separately from the two southernmost sampling locations in Michigan. When further structuring was explored, the two MI locations (AD-MI and SBD-MI) separated out into their own genetic clusters, though overlap was clearly visible when viewing discriminant functions. The two more southern sampling locations in Michigan also had higher levels of genetic diversity than the other five sampling locations.

There are likely multiple factors contributing to the genetic patterns that we observe across these populations of invasive baby’s breath. The increased levels of genetic diversity observed in the SBD-MI and AD-MI populations compared with the other sampled locations could be due to a combination of population size and connectivity. Populations located in SBD-MI tend to be much larger than other locations sampled in this study. Larger populations tend to be more robust to the effects genetic drift and can help resist the effects of inbreeding, helping to retain diversity within these populations (see Ellstrand & Elam, 2003). Another possible reason for the patterns found here is that sampling locations spread across the western U.S. are more isolated than the two southernmost Michigan locations, which may be contributing to lower levels of genetic diversity among these areas. Several sample locations (CH-WA, OT-MN) occur in relatively fragmented or space-limited environments, which may result in a lack of gene flow to other populations of *G. paniculata* growing nearby or prevent its spread altogether. The close geographic proximity between SBD-MI and AD-MI could also be maintaining some gene flow between these populations. However, many of our other sample locations with limited genetic diversity (OB-WA, PS-MI, KR-ND) were part of a contiguous landscape that was not obviously limiting to expansion.

One potential explanation for the distinct genetic clustering we observed with our data is that the populations of SBD-MI and AD-MI that were established in the 1940’s could have been founded by individuals from the existing PS-MI population. SBD-MI and AD-MI could then have significantly diverged from the initial source over the past 50 years. However, this scenario seems unlikely. Our data show that SBD-MI and AD-MI have higher levels of genetic variation compared to PS-MI and a number of private alleles were found in both SBD-MI and AD-MI that are not present in PS-MI. Additionally, chloroplast microsatellite data from a previous study (Leimbach-Maus et al., 2018a) show that the SBD-MI and AD-MI populations have distinct DNA haplotypes compared to the PS-MI population and other more northern Michigan populations not included in this study. The combination of these data suggest that SBD-MI and AD-MI are likely not the result of serial founding events from the source population of PS-MI.

A more likely explanation for the distinct patterns observed among our populations could be a signature of *G. paniculata’*s horticultural past. The earliest occurrences of *G. paniculata* populations across several different regions in the U.S. coincides with its initial introduction to N. America though seed sales. Based upon seed catalogs from the Biodiversity Heritage Library, *G. paniculata* was promoted as a garden ornamental as early as 1856 in the *Farmer’s Promotion Book* (Reinhold, 1856). By 1868 at least two seed distributors (J.M. Thorburn & Co, NY and Hovey & Nichols, Chicago) were selling *G. paniculata* in their catalogs in New York and Chicago; the earliest herbarium records of *G. paniculata* collected in the United States were from CA (1907), MN (1896), MI (1913), and NY (1894) (Table S2). We hypothesize that when *G. paniculata* initially invaded N. America in the late 1890’s there may have been little standing genetic diversity present in the garden cultivars being grown at the time. Additionally, the number of oversees distributors of seeds may have been further restricting possible diversity. These potential limitations to genetic diversity during the early periods of invasion are likely why some of our populations cluster together, despite the large geographic distances between them. According to herbarium records, populations of *G. paniculata* in SBD-MI and AD-MI were not established until the later 1940’s, when *G. paniculata* had become a more popular garden ornamental. This increased popularity likely led to the number of seed distributors being greatly increased. We suggest then that the genetic patterns observed in this study among populations of *G. paniculata* are a signature of the horticultural past that helped facilitate its invasion into N. America.

One confounding factor to our genetic analyses is that tissue from the SBD-MI and AD-MI populations was collected two-years prior to the other locations (2016 compared to 2018), which could potentially impact our structure results. However, the same Leimbach-Maus et al. (2018a) study examining the genetic structure of population throughout west Michigan also found that baby’s breath populations north of the Leelanau Peninsula (i.e., PS-MI) group in a cluster that is distinct from both SBD-MI and AD-MI. Samples from this study were all collected in the same year. Thus, the distinct clustering of PS-MI from the SBD-MI and AD-MI populations appears to be well supported.

Invasion curves created at multiple geospatial scales help assess the current invasion status of *G. paniculata* across its introduced range in North America. Herbarium records compiled for North America indicate that *G. paniculata* has likely not yet reached a plateau phase, and its range could still be expanding. When this larger invasion is viewed at a finer geospatial scale, additional trends become visible. Herbarium records collected from the geographic area of cluster 1 (Washington, North Dakota, Minnesota, and northwestern Michigan) show a lag period that ended in the 1920’s as *G. paniculata* collection increased in new localities and its range began expanding. The invasion curve created for cluster 2 (Michigan south of the Leelanau Peninsula) shows that the expansion phase was already in process during the first collection period or shortly after, with little lag phase observed. Whether this is because *G. paniculata* was present within the region prior to this period but not collected until the 1940’s, or whether populations were not present in this area until the 1940’s and began spreading rapidly shortly after introduction is unclear. Regardless, the expansion in this region was in progress in the mid 1940’s, with a plateau in new localities invaded taking place around 1970. These distinct expansion phases could suggest at least two separate periods of invasion occurring across our sampled range, one expanding in the 1920’s and another in the 1940’s.

This combination of genetic and herbarium data offers valuable insight into the invasion of a problematic weed across a large portion of its invaded range. Using genetic analyses, we were able to infer the likely number of distinct invasion events across a large geographic spread of invasive weed populations. Using the data gleaned from these analyses, we were then able to construct informed invasion curves that reveal trends that would otherwise have been obscured in the large pool of available data. This combination of genetic analyses as *a priori* information for the construction of herbarium-derived invasion curves proves a powerful method for extracting information on the invasion status of distinct invasion events, as well as maximizes the benefits of freely available data. In an era of increased invasions and dwindling conservation funding, the use of existing data in the most effective and informed way possible is paramount for the continued effective management of invasive species and increased understanding of invasion success.

In conclusion, this study offered insight to the population structure and invasion status of a *Gypsophila paniculata* in its introduced N. American range. Our data suggest that the distinct population clusters observed through genetic analyses are likely explained by the species’ history as a horticultural species, a characteristic that facilitated its spread to the continent. When viewed in light of these genetic clusters, herbarium data further supported the presence of at least two invasion events, evidenced by unique expansion phases across the species’ range. Combining herbarium records with genetic analyses has provided a more complete analysis of the invasion history of this species, and this type of work would serve as a useful tool for characterizing the invasion status of other invasive populations.

## Acknowledgements

The authors would like Hailee Leimbach-Maus and Emma Rice for their help in the field. Additionally, the authors would like to thank the Environmental Protection Agency – Great Lakes Restoration Initiative (C.G.P., Grant #00E01934), the Michigan Botanical Foundation, and Grand Valley State University for financial support.

## Biosketches

- Sarah K. Lamar’s research interests rest at the intersection of molecular ecology and conservation biology, using genetic techniques to inform management and practical conservation. Her particular interests are related to 1) adaptation to environmental changes and 2) the genetic bases of behavior.
- Charlyn G. Partridge’s work encompasses both field and laboratory studies and employs a variety of behavioral, molecular, and genetic techniques to address questions related to: invasive species management, sexual selection, and alternative mating strategies.

## Supporting/Supplemental Information

**Figure S1.**
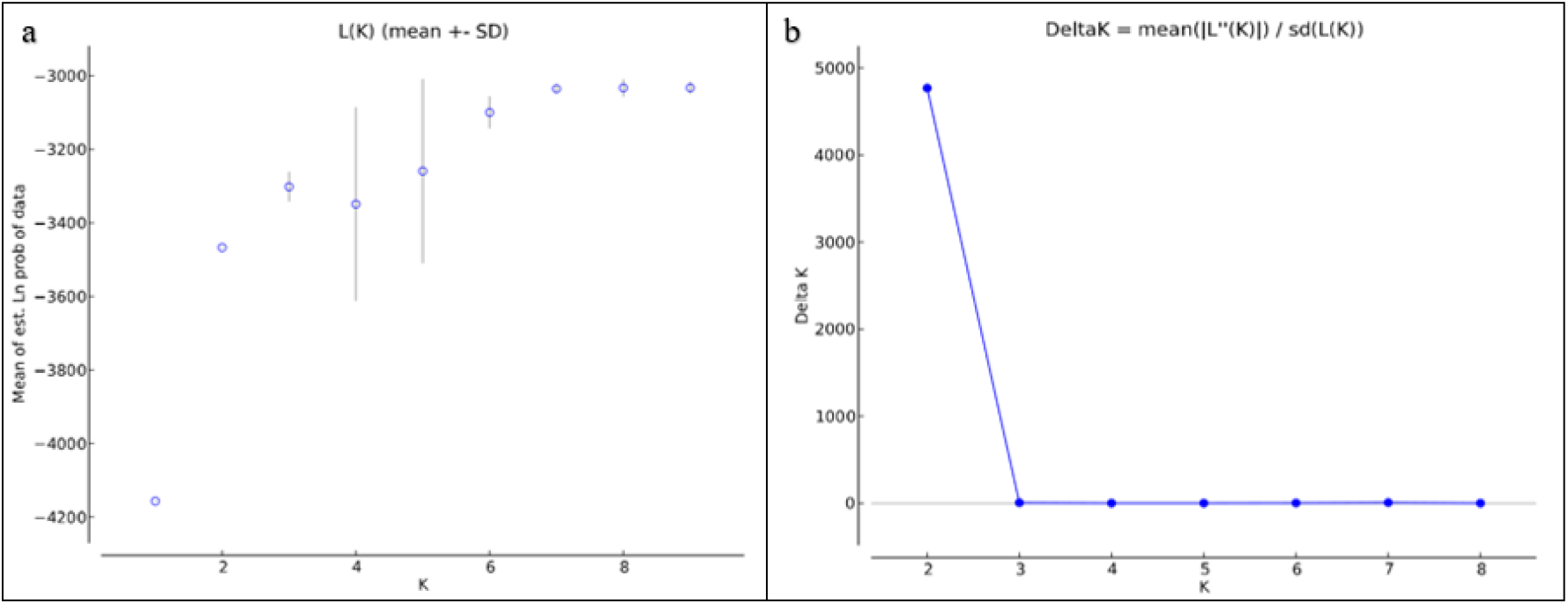
Bayesian cluster analysis of seven sampling locations of baby’s breath (G. *paniculata)* genotyped at 14 microsatellite loci, gathered from the program STRUCTURE (Pritchard et al. 2000). (a) Mean L(K) (±SD) over 10 runs for each value of K (1-9). (b) Evanno’s ΔK (Evanno et al., 2005) where the highest rate of change indicates the highest likelihood of cluster numbers. This analysis was conducted without prior sampling location information. Two genetic clusters were inferred.

**Figure S2.**
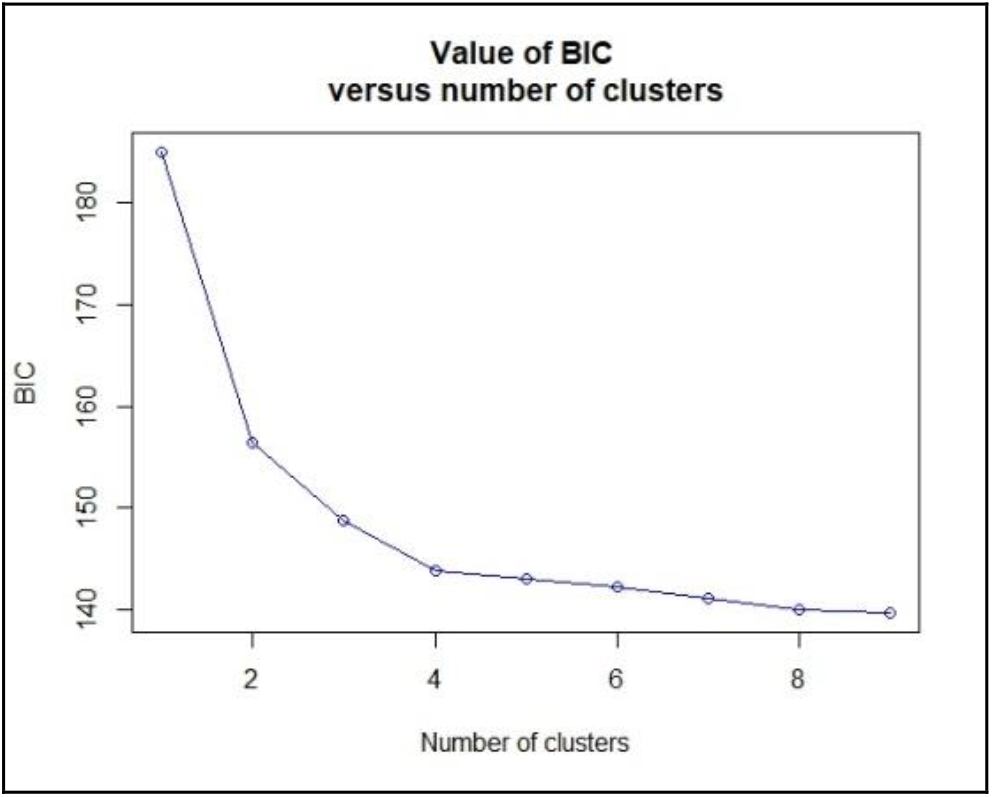
Bayesian Information Criterion for a DAPC of seven sampling locations of baby’s breath *(G. paniculata)* genotyped at 14 microsatellite loci, created using the package *‘adegenet’* in R (Jombart & Collins, 2015; Jombart et al., 2010). The inflection point suggests the supported amount of genetic clusters present; both a *K* of 2 and 3 were considered in analysis.

**Table S1.**
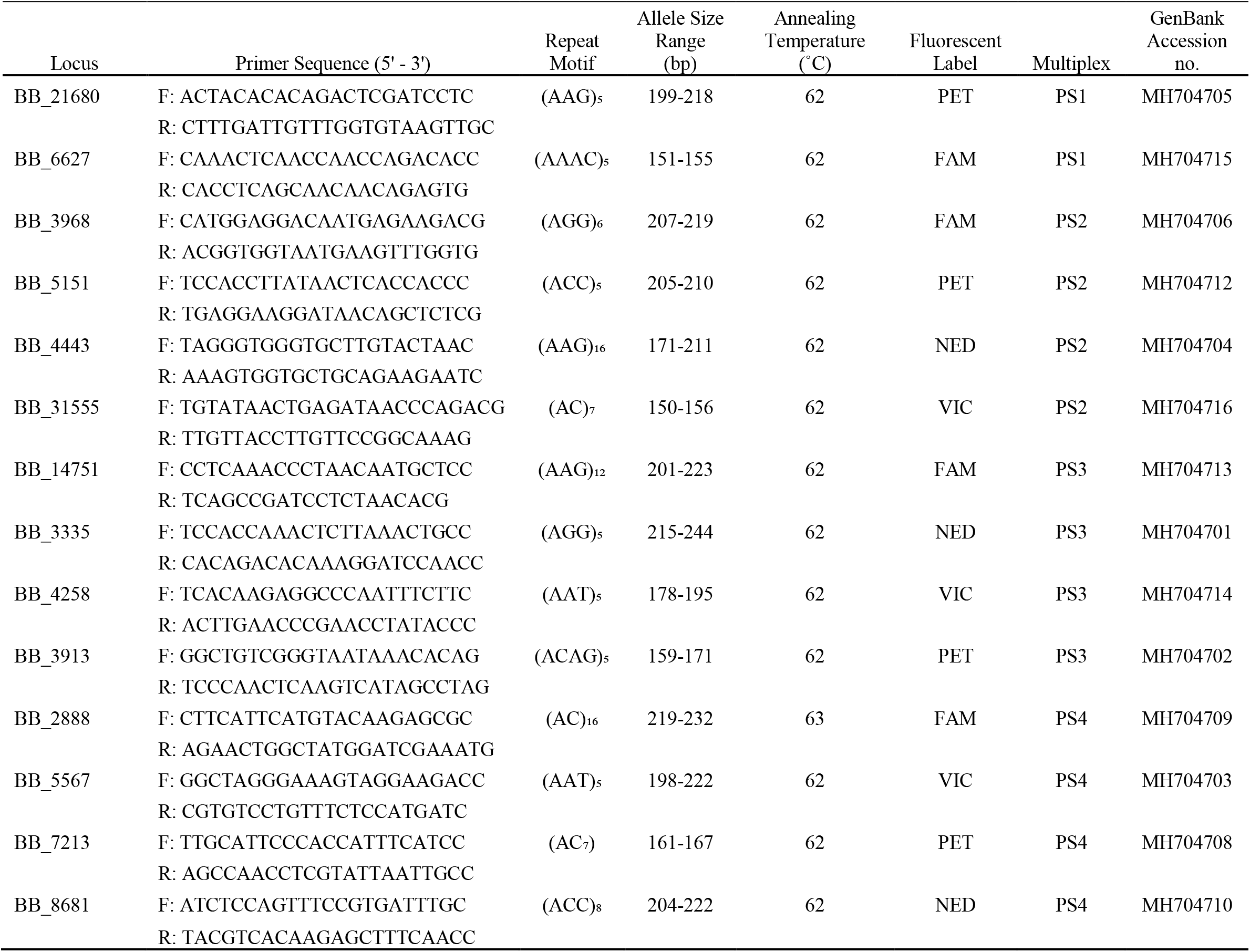
Details for 14 microsatellite (nSSR) loci specific to *G. paniculata* developed by Leimbach-Maus et al. (2018b) and used in this study.

**Table S2.**
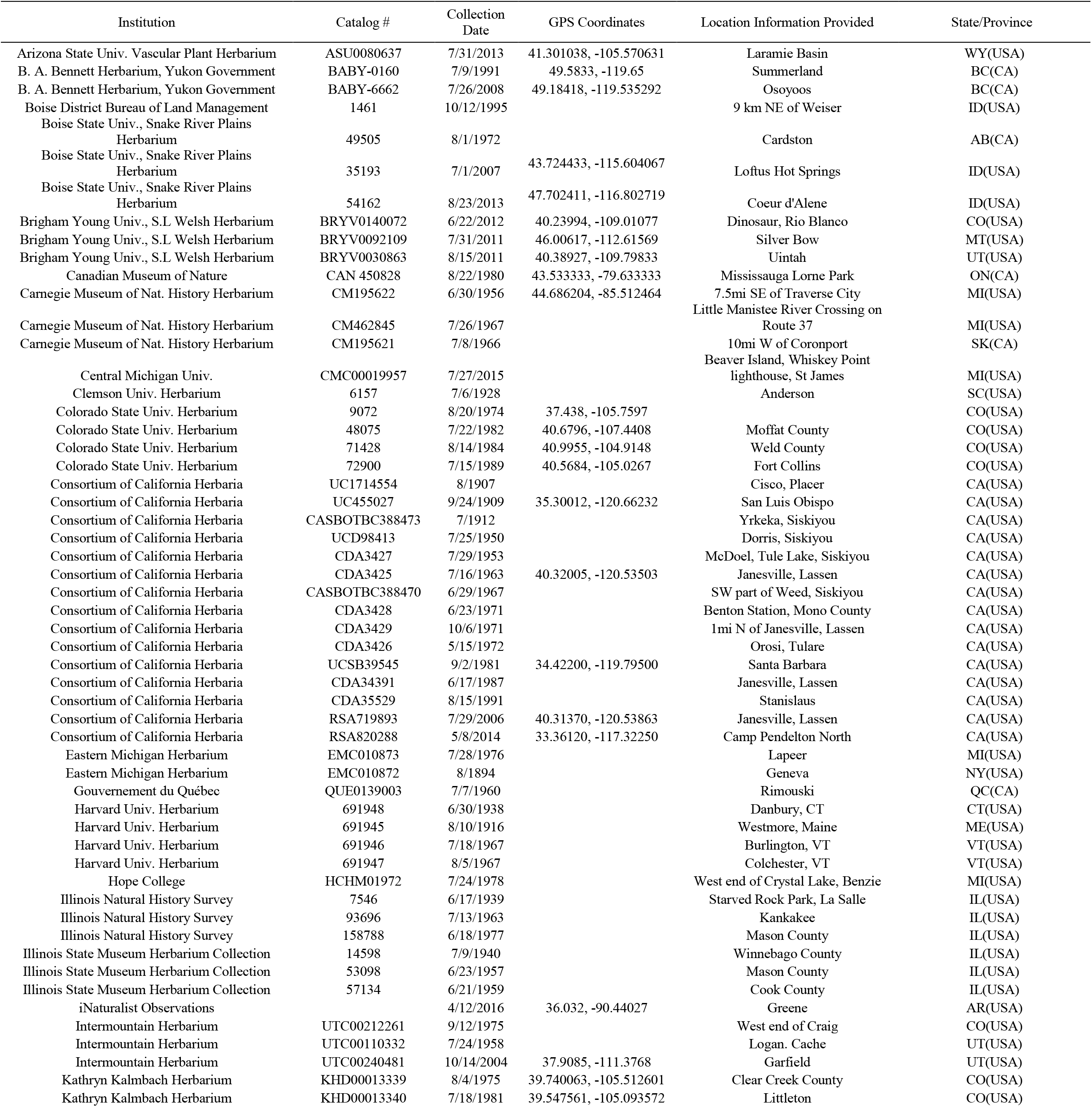
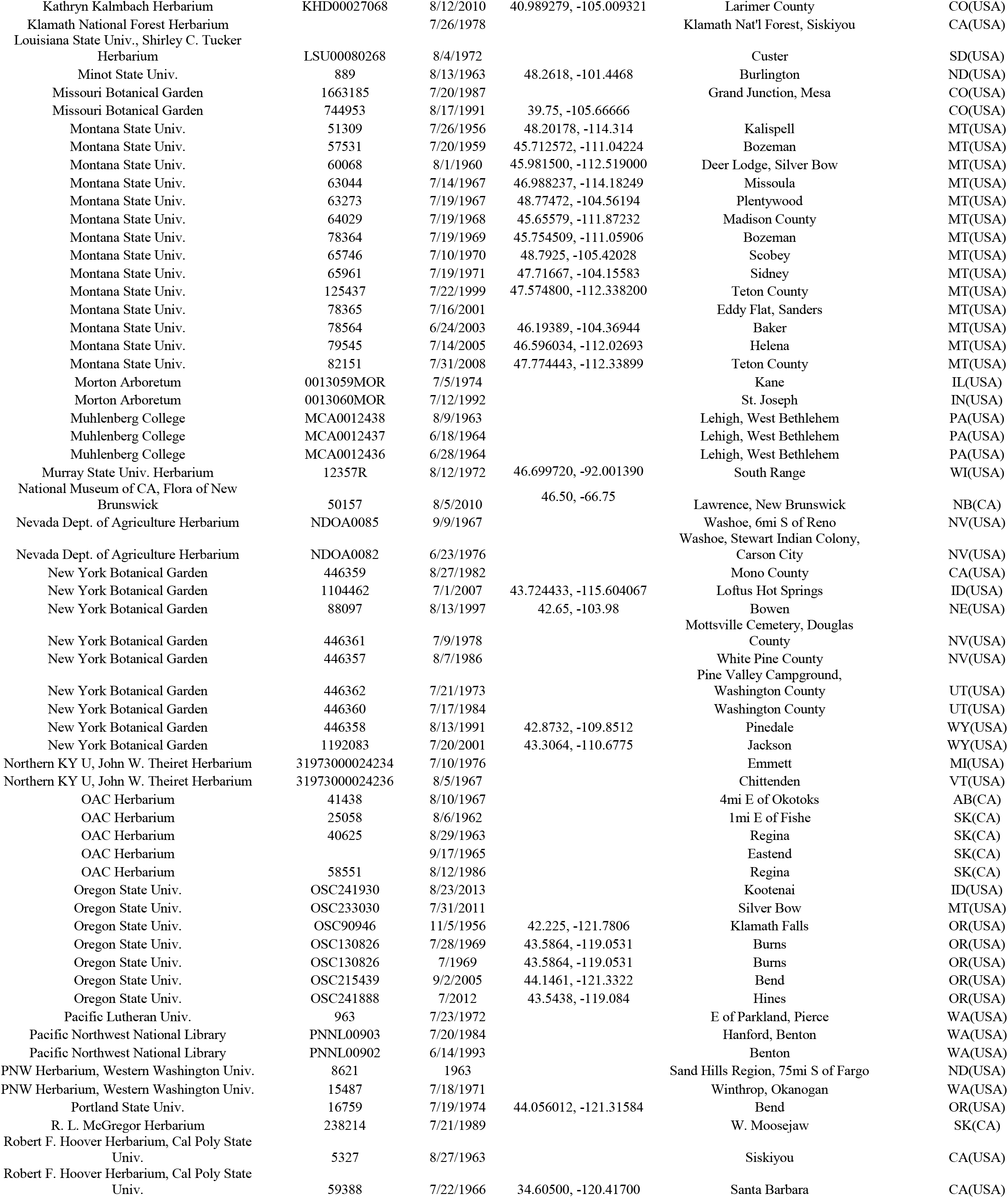
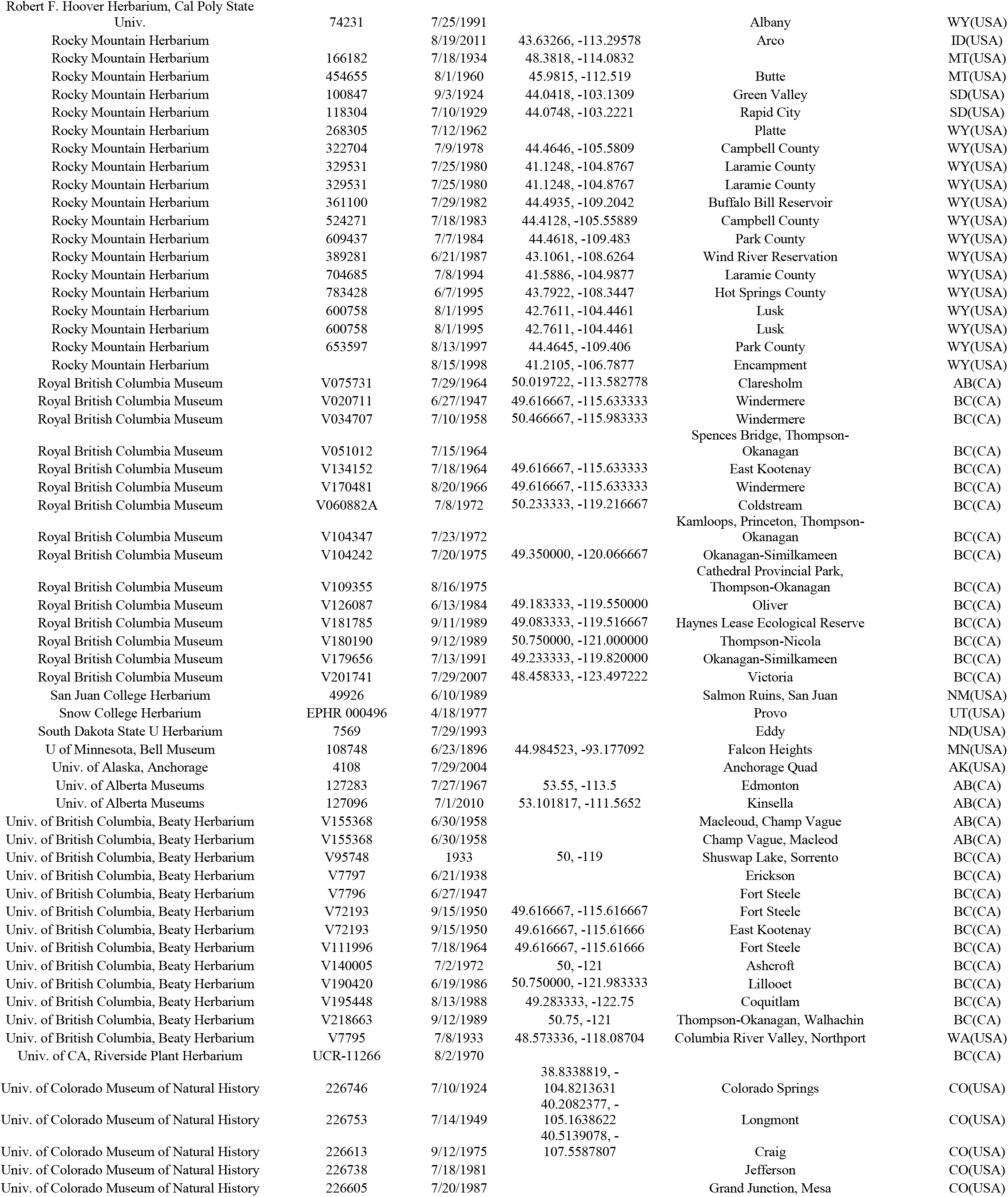
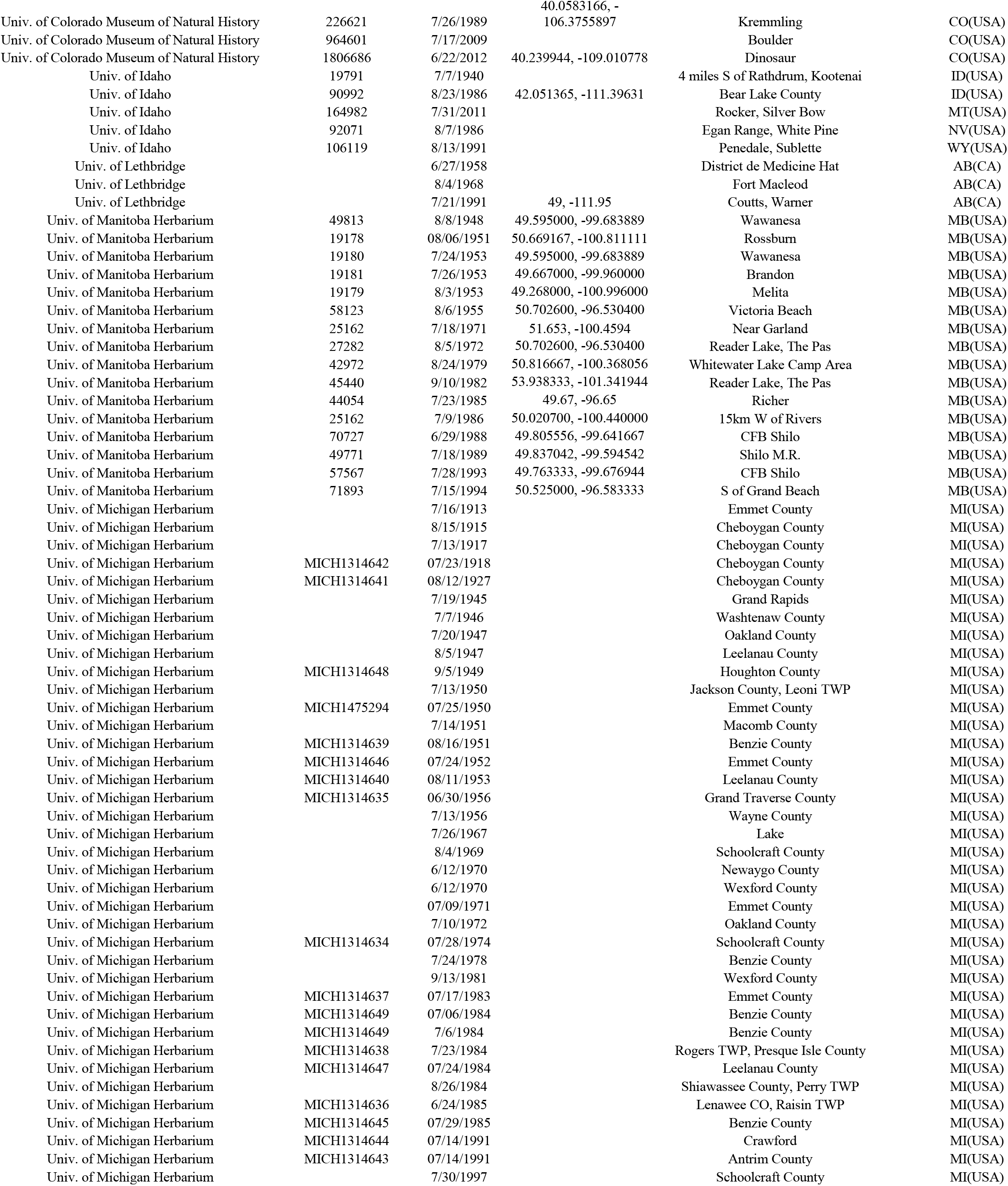
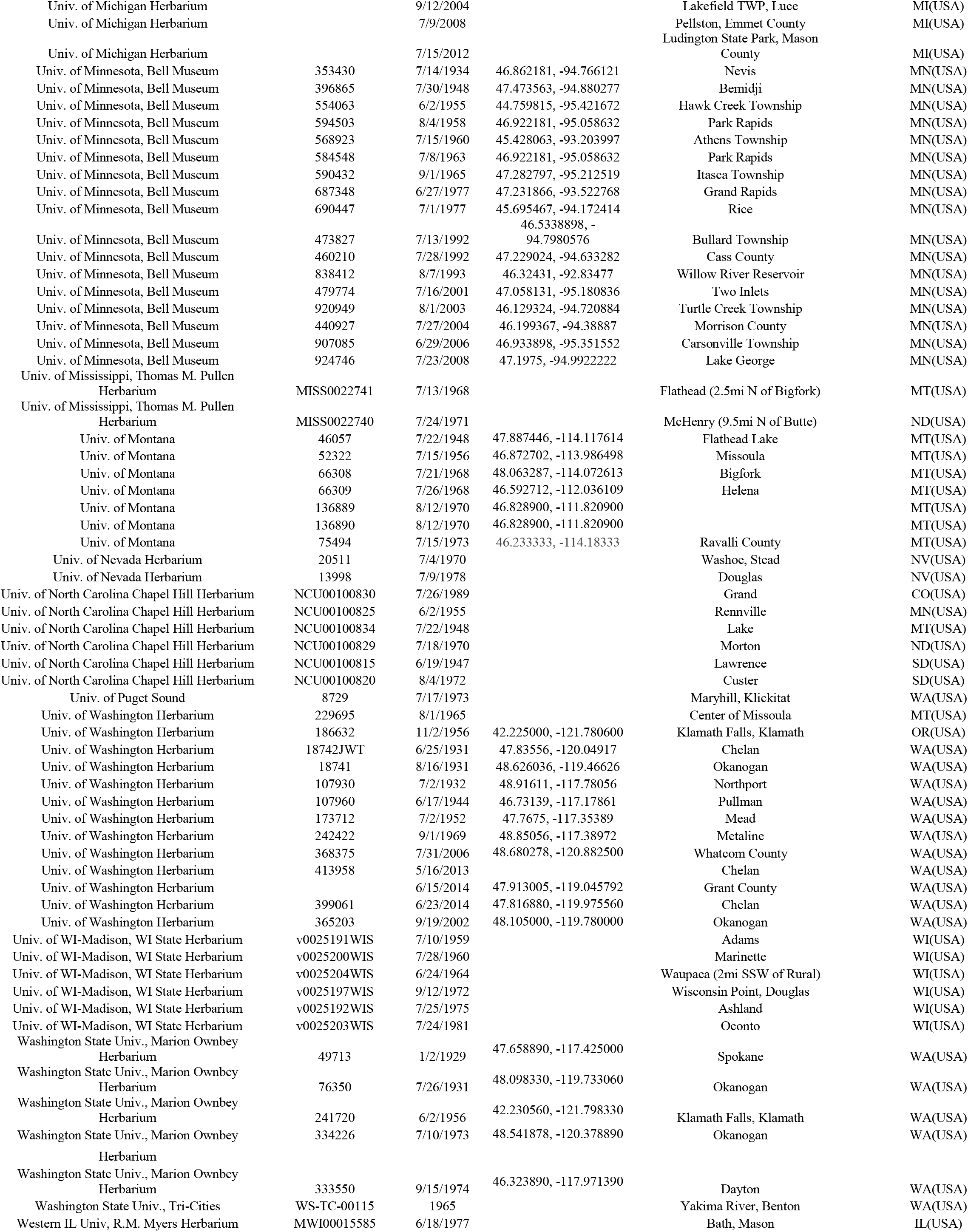
Details for *G. paniculata* herbarium records used in this study.

